# Multiplatform lipid analysis of the brain of aging mice by mass spectrometry

**DOI:** 10.1101/2024.09.25.614823

**Authors:** Punyatoya Panda, Christina R. Ferreira, Bruce R. Cooper, Allison J. Schaser, Uma K. Aryal

**Affiliations:** Department of Comparative Pathobiology, Purdue University, West Lafayette, IN 47907, USA; Bindley Bioscience Center, Purdue University, West Lafayette, IN 47907, USA; Department of Speech, Language, and Hearing Sciences, Purdue University

**Keywords:** Aging, DESI imaging, proteomics, targeted lipidomics, untargeted lipidomics

## Abstract

Lipids are an integral part of brain structure and function and represent about 50% of the dry weight of the brain. Despite their importance, the complexity and variations in the abundance of brain lipids due to aging remain poorly understood. For maximum coverage and multi-platform validation, we applied three complementary mass spectrometry-based analytical approaches: multiple reaction monitoring (MRM) profiling, untargeted liquid chromatography tandem mass spectrometry (LC-MS/MS), and desorption electrospray ionization-MS imaging (DESI-MSI). We used three different age groups of mice, namely adult (3-4 months), middle-aged (10 months) and old (19-21 months). Phospholipids including phosphatidylcholine (PC), phosphatidylethanolamine (PE) and phosphatidylglycerol (PG) showed higher abundance, while phosphatidylinositols (PI) and phosphatidylserines (PS) generally showed lower abundance in the brains of old mice compared to adults or middle-aged mice. Polyunsaturated fatty acids, such as docosahexaenoic acid (DHA) and arachidonic acid (AA), as well as hexosylceramides (HexCer), sulfated hexosylceramides (SHexCer) and sphingomyelins (SM) were among the most abundant lipid species in the brains of old mice. DESI-MSI showed variations in the spatial distribution of many of the lipids confirmed by MRM and LC-MS/MS profiling. Interrogation of lipidomic data with recent proteomics data obtained from the same tissues revealed changes in the abundance and phosphorylation levels of several proteins potentially linked to ceramide (Cer), hexosylceramide (HexCer), fatty acids (FA), phosphatidylinositol (PI), sphingomyelin (SM) and sulfatides (SHexCer) metabolism and correlated well with the multiplatform lipid surveillance. Our findings offer insight into age-dependent changes in brain lipid profiles and their potential contribution to age-related cognitive decline.

## Introduction

Lipids are an integral part of many biological systems such as cell structure, energy storage, signaling, and metabolism [1, 2]. The presence of multiple acyl chains with varying lengths, degree of saturation, and *cis-trans* configuration contribute to lipid complexity and functional specificity [3, 4]. Phospholipids, glycerolipids, sphingolipids, and sterols have different functions and each lipid class has its own characteristic structure and function, further contributing to the general complexity and analytical challenges of understanding the role of lipids in healthy aging and disease [3, 5, 6]. Recent advances in mass spectrometry (MS) techniques are attracting unprecedented interest in the analysis of lipids because MS is a powerful technique that allows for the most detailed characterization of lipid structure and function [7]. The MS analysis of lipids is performed mainly by either direct infusion or LC-based tandem MS (MS/MS). Direct infusion coupled with triple-quadrupole (QqQ) MS is quick, easy, and more sensitive [19], but the interpretation of MS/MS spectra is also difficult due to the high degree of overlaping signals [20–22]. LC separation prior to MS/MS analysis can solve this problem by reducing sample complexity and chromatographic separation of overlapping peaks. However, the lack of suitable internal standards and the limitation of existing MS/MS database libraries [21] hinder the identification of many peaks observed using untargeted LC-MS/MS. Improving existing MS/MS database libraries and utilizing complementary techniques can help address these limitations. The Cooks group at Purdue University has recently developed a shotgun MRM-based profiling method, which is based on known functional groups rather than individual lipid molecular species [23, 24]. This shotgun MRM workflow does not utilize LC and relies on specific ion transitions corresponding to the functional head group and fatty acid composition. This method is particularly useful if characteristic headgroup fragments or neutral losses are available, as is the case for most phospholipids [25, 26] and sphingolipids [27, 28]. The selection of MRM scans is based on the well-known precursor ion scan (Prec) and neutral loss scan (NL) and a priori knowledge of lipid fragmentation both from experiments and from previous literature, primarily from LipidMaps [23, 24]. In addition to mass spectrometry lipid analysis methods based on homogenized tissues, there is the possibility of spatially determining the location of lipids. The DESI imaging technique is a well-established mass spectrometry imaging method to obtain spatial information on the distribution of lipids within tissues. The nondestructive nature of this technique combined with the enhanced spatial resolution and versatility of current DESI-MSI techniques in lipid detection are driving advances in our understanding of lipids and their implications in health and disease [8]. As the technique continues to evolve, it is likely to play an increasingly important role in aging research.

Like proteins, lipids also have extensive cellular functions [9] and play a critical role in biological aging. Aging is a major risk factor for the development of neurodegenerative diseases such as Alzheimer’s (AD) and Parkinson’s (PD) [10–12] . The brain has the second highest lipid content after adipose tissues, accounting for about 50% of its dry weight [13]. However, changes in lipid composition and abundance in the brain due to aging are not fully understood. Brain lipids consist mainly of cholesterol, phospholipids, such as phosphatidylcholine (PC) and phosphatidylethanolamine (PE), and sphingolipids [14, 15]. Any changes in lipid composition, spatial distribution, saturation, and altered ratio can influence bioenergetics, membrane integrity, and signaling [16] which declines with aging. Abnormal accumulation or disruption of lipid metabolism within the brain contributes to neurodegenerative disorders [17, 18]. Advances in the field of MS have enabled us to identify new lipids with temporal and spatial resolution [19–21], however, characterizing complex lipidomes at the individual species level remains challenging.

Therefore, despite our growing understanding of brain lipids, knowledge of their composition, spatial organization, structure, and function in aging brains is still limited. In this study, we applied MRM profiling, LC-MS/MS-based untargeted lipidomics, and DESI-MSI to determine differences in the abundance profiles and spatial distributions of lipids. MRM profiling, we selected 3,246 MRM transitions comprising phospholipids, triacylglycerols, diacylglycerols, ceramides and acylcarnitines across the brains of 13 mice divided into three age groups. We considered 576 MRMs detected, including phosphatidylcholines (PCs), phosphatidylethanolamines (PEs), and free fatty acids (FFAs) among the most abundant classes of lipids in the brain. The non-targeted LC-MS/MS resulted in the identification of 226 lipid species, most of which were also identified in the MRM profiling workflow. Using DESI-MS imaging, we were able to observe the spatial distribution of several of these lipids and identify age-dependent changes in brain lipids, which may contribute to age-related neurodegenerative diseases.

## Materials and Methods

### Chemicals

Methanol, chloroform, and acetonitrile were purchased from Fisher Scientific. The bicinchoninic assay kit (BCA) and formic acid were purchased from Thermo Fisher Scientific. Tetraethylammonium bromide (TEAB) and bovine serum albumin (BSA) were purchased from Sigma.

### Animal model

The mice were housed by Purdue’s Laboratory Animal Program in a light-dark cycle and temperature and humidity-controlled vivarium and maintained on an ad libitum food and water diet. The experiments were carried out according to the relevant guidelines approved by the Purdue Institutional Animal Care and Use Committee (IACUC) described in the protocol 2008002069. Every effort was made to minimize the number of mice used and their suffering. All animals used in this study were wild-type male C57BL/6 littermates from the A53T SynGFP mouse line [22] which is currently maintained at Purdue University.

### Tissue Collection and Homogenization

Mice were anesthetized with 4% isoflurane in 100% oxygen at a flow rate of 3 L / h in a surgical box. During anesthesia, the mice underwent rapid decapitation and tissues were dissected. Following dissection, the brain was removed and washed by placing it in a 50 ml falcon tube containing 1x PBS buffer on ice and immediately flash frozen in liquid nitrogen and stored at - 80°C until further use.

Whole brains were collected from five young adult (3-4 months), which are referred to as adult hereafter, three middle-aged (9-12 months) and five old (19-21 months) male mice. The entire brain tissues were cut and shredded with the Shredder SG3 kit (Pressure Biosciences, Inc., MA) in 50 mM TEAB in deionized water supplemented with protease and phosphatase inhibitors. Phenylmethylsulfonyl fluoride (PMSF) was used as a serine protease inhibitor to a final concentration of 1 mM. A mixture of sodium fluoride (NaF), Na-molybdate (Na_2_MoO_4_), Na-orthovanadate (Na_3_VO_4_), and Na-beta glycerophosphate (C_3_H_19_Na_2_O_11_P) at a final concentration of 10 mM was used as phosphatase inhibitors. The ground tissues were transferred to prefilled Precellys soft tissue homogenizing CK14 tubes (Bertin Corp., Rockville, Maryland, USA) and homogenized six times in a hard tissue homogenizer at 3200 rpm for 3 x 20 s in each cycle. The protein concentration was measured using a BCA protein assay kit with BSA as a standard. Lysate volumes equivalent to 100 µg of protein were taken for each sample for lipid extraction, thus normalizing the amount of lipid extracted by the same protein concentration.

### Lipid Extraction

Lipopolysaccharide extraction was performed by Bligh-Dyer method [23]. Brain lysates containing 100 µg of protein (equivalent volume) were placed in a microcentrifuge tube and made up to 200 µL with 1x PBS buffer. Then 500 µL of chloroform was added and briefly mixed, followed by 400 µL of methanol to make a one-phase solution. The solution was kept at 4 ° C for 10 min, and 250 µL of water was added and vortexed, followed by centrifugation at 4000 xg for 10 min. The top phase consists of the polar metabolites, the middle layer consists of proteins, and the bottom phase consists of lipids. The top phase was carefully removed and then the bottom phase was carefully transferred to a new tube without removing protein from the intermediate layer and dried without heat in a vacuum centrifuge (Vacufuge Plus, Eppendorf, Enfield, CT) and stored at -80°C until MS analysis.

### MRM profiling analysis of lipids

The dried lipid extracts were resuspended in 200 µL of acetonitrile: MeOH: 300 mM NH_4_Ac (3:6.65:0.35 v/v/v) to obtain a stock solution. Two µL of the stock solution of each sample was diluted 100 times by adding 198 µL the same solvent as the stock solution to achieve the desired concentration range. Eight µL of diluted samples were introduced into an Agilent 6410 triple quadrupole (QQQ) MS (Santa Clara, CA) through a G1367A 1100 series autosampler (Santa Clara, CA) at a flow rate of 10 µL/min and a pressure of 100 bar. For MRM profiling analysis, samples were electrosprayed directly into the QQQ without chromatographic separation with a capillary voltage of 4 kV for the positive ion mode and 3.5 kV for the negative ion mode, 25 ms dwell time, gas flow at 5.1 L/min at 300, temperature at the ESI source gas temperature 300°C [24]. The collision energies as reported in Supplemental Dataset 1 Table S1. Acetonitrile with 0.1% formic acid (FA) was used as a solvent between sample injections. The collision energies reported by Reis *et. al,* 2023 [25]. MRM profiling analysis was performed by screening the samples for specific ion transitions corresponding to different lipid classes and fatty acid composition based on an a priori knowledgebase or LIPID MAPS database. Data acquisition was performed using multiple sample injections to interrogate samples for MRM related to different lipid classes, as described by Reis *et al.,* 2023. Each sample was screened for 3,246 MRMs that span 11 different classes of lipids [25], with separate injections for each type of MRM method for each sample.

### Untargeted LC-MS/MS profiling of lipids

Dried lipid extracts prepared as described above were resuspended in 50 µL of ACN: MeOH: H_2_O (3:5:2, v/v/v) and 5 µL were injected into the LC column. LC separation was performed on an Agilent 1290 system (Palo Alto, CA), with a flow rate of 0.40 ml / min. Lipoproteins were separated using a Waters BEH C18 column (1.7 um, 2.1 x 100 mm) (Milford, MA), where mobile phase A was 10 mM ammonium acetate and 0.1% formic acid in ddH_2_O and B was 10 mM ammonium acetate and 0.1% formic acid in acetonitrile (HPLC grade). The initial conditions were 65:35 A:B, held for 0.5 min, followed by a linear gradient to 20:80 at 5 min, then 5:95 at 10 min with an isocratic hold until 15 min. Column equilibration was performed by returning to 65:35 A:B at 17 min and holding until 21 min. The mass analysis was obtained in either positive or negative ionization mode using an Agilent 6545 Q-TOF mass spectrometer with ESI capillary voltage 3.5 kV. MS/MS was performed in Data-Dependent Analysis (DDA) mode, with a range of 100 –1200 *m/z* and fixed collision energies of 10, 20, and 40 eV. The mass accuracy was monitored by infusing Agilent reference mass standards. Data acquisition was performed using Mass Hunter Workstation Data Acquisition software (Agilent), and the data was qualitatively analyzed using Qualitative Analysis 10.0 (Agilent). Peak deconvolution, integration, and statistical analysis were performed using MS-DIAL (v. 4.9.221218). Peak annotations were performed using the MassBank of North America MS/MS library, based on authentic standards (v. 16). The mass tolerances for identification were 0.005 Da for MS1 and 0.01 Da for MS2.

### Data Processing

#### Shotgun MRM Profiling

Data processing was carried out using an internal script that summed the ion intensity of each MRM over the acquisiton time, and the data was filtered so that only MRMs with intensities greater than 30% of the blank measurements were considered ’detected’ for the profiling analysis. Data normalization was done by the sum of the intensities for each class, excluding the internal standard data for each class separately. The detected normalized lipids from each class were combined into a single list. Statistical analysis was performed using Perseus (version 2.0.7.0) and OriginPro (Version 2022, OriginLab Corporation, Northampton, MA, USA). Normalized relative ion intensities were multiplied by a factor of 10 8 and then log2-transformed. Multiple sample ANOVA test was performed, with a 5% FDR (False Discovery Rate). Principal component analysis was performed to visualize the variation within the three groups (adult, middle, and old). The association of the different classes of lipids was evaluated using Pearson’s correlation. Lipids with *P<0.05* across the three ages were considered significantly changing lipids and were further considered for biological interpretation.

#### Untargeted LC-MS/MS

MS-DIAL (v. 4.9.221218) was used to process LC-MS / MS data including spectral deconvolution, peak identification, and data processing. A retention time window of 0.2-15.5 min was considered for the data processing steps. Statistical analysis was performed with Perseus (version 2.0.7.0) [26] and OriginPro (Ver. 2022, OriginLab Corporation, Northampton, MA, USA). The intensities were log_2_ transformed and used for statistical analysis using multiple sample ANOVA tests. Data acquired in positive- and negative-ion mode were analyzed separately. From the list of significantly changing lipids, only lipids that had MS1, MS2 information and could be matched to the MSP spectral database or the mass spectral atlas database were considered confident identifications. These significantly changing lipids with *a value p* <0.05 from both positive and negative ion modes were combined into a single sheet for downstream analysis and biological interpretation.

The lipids identified by the targeted and untargeted methods were compared considering the lipid classes that were detected in both methods (phospholipids, acylcarnitine, ceramides, and sphingomyelins, and di- and triacylglycerols (DG and TG), etc. Specific lipids that were significantly changing among the three age groups of the untargeted data and mapped to the same classes of lipids from the targeted data were highly confident detection and were considered for further analyses and biological interpretation.

### DESI-MS Imaging

For DESI-MSI, a subset of brain tissues from three age groups of mice were embedded in carboxymethylcellulose (CMC) and stored at -80°C. The frozen tissues were cut into sections in a Leica CM1860 cryostat at -10 ° C with a thickness of 10 µm. The CMC embedding medium was prepared using a protocol described before [27]. Briefly, 10g of sodium carboxymethylcellulose was heated with 450 ml of deionized water and incubated overnight at 70 ° C with stirring and brought to 500 ml with deionized water. It was cooled and refrigerated until further use.

The sagittal tissue sections from these different age groups were placed side by side on the same slide for imaging. However, different slides were used for positive and negative ion modes. DESI imaging was performed on a Synapt XS mass spectrometer connected to a XS DESI source (Waters Corporation, Milford, MA). Frozen tissue sections were dried at room temperature before use. Mass spectra were collected by running a series of scans across the tissue sections using ambient ionization, and data were collected using Mass Lynx v. 4.2. with m / z 100-1,200 as mass range for the positive ion mode and *m/z* 50-1200 in the negative ion mode at 150 μm pixel resolution. MeOH: H_2_O 98:2 (v/v) was used as the solvent and sprayed at a rate of 3 µl/sec. The source temperature was 150 ° C. The pentapeptide leu-enkephalin (Tyr-Gly-Gly-Phe-Leu) was used as the mass-to-mass lock correction using *m/z* 556.2771(+) and 554.2615(-) as references. For the negative ion mode, Leu-Enk was used at a concentration of 0.03%, and for the positive ion mode, it was doubled to 0.06% to reach the needed ion intensities. Before viewing results, the data sets were subjected to continuous lock mass correction (CLMC) by leucine enkephalin. Next, high-definition imaging (HDI) software was used to process and convert the MS data into the ion imaging format. Signal intensities were normalized to the total ion current in the HDI software. Annotations and segmentation were performed using Waters Microapps (MSI Analyte browser, MSI Segmentation 2.1.2) using the corrected lock mass calibration (CLMC) file for both positive and negative modes, respectively. Lipid annotation was also manually curated using HMDB, Lipid Maps and literature reporting DESI-MS imaging of brain tissue, especially including product ion scans [28]. The 5,000 most intense ions were acquired in HDI software to visualize at an MS resolution of 15,000.

The corresponding spectra for the lipids of interest were viewed in the MassLynx Spectra viewer. Specific regions of interest (ROIs) were designated in images of whole brain tissue sections in the HDI software. Following analyte clustering in MSI segmentation and UMAP classification, ROIs were designated for six distinct regions of the brain that were clearly separated in the sagittal sections, namely cerebellum, cerebral cortex, midbrain, pons and medulla, thalamus, hypothalamus, and basal ganglia, and their spectra were compared to draw insights about specific lipids distributed only in those regions and highly abundant lipids in those regions.

## Pathway Analysis

Metscape was used as an app plug-in in Cytoscape to map the pathways represented by the identified lipids and for linking genes to lipids [29, 30]. The KEGG IDs were retrieved for the significantly changing lipids as input in Metscape. We also compared current lipidomics data with proteomics and phosphoproteomics data that we collected from the same animals (unpublished). For this comparison, only proteins that were identified as significantly different between adult and old mice and are known to be related to significantly changing lipids were considered.

## RESULTS

### MRM profiling of different lipid classes in brains of aging mice

Figure 1 provides an overview of the experimental and bioinformatic workflow that combines three different analytical methods. The MRM profiling is LC-free since it uses flow injection, and it is based on the characteristic head group fragments or neutral losses detected as product ions of an MRM (or ion transition).

**Figure 1:**
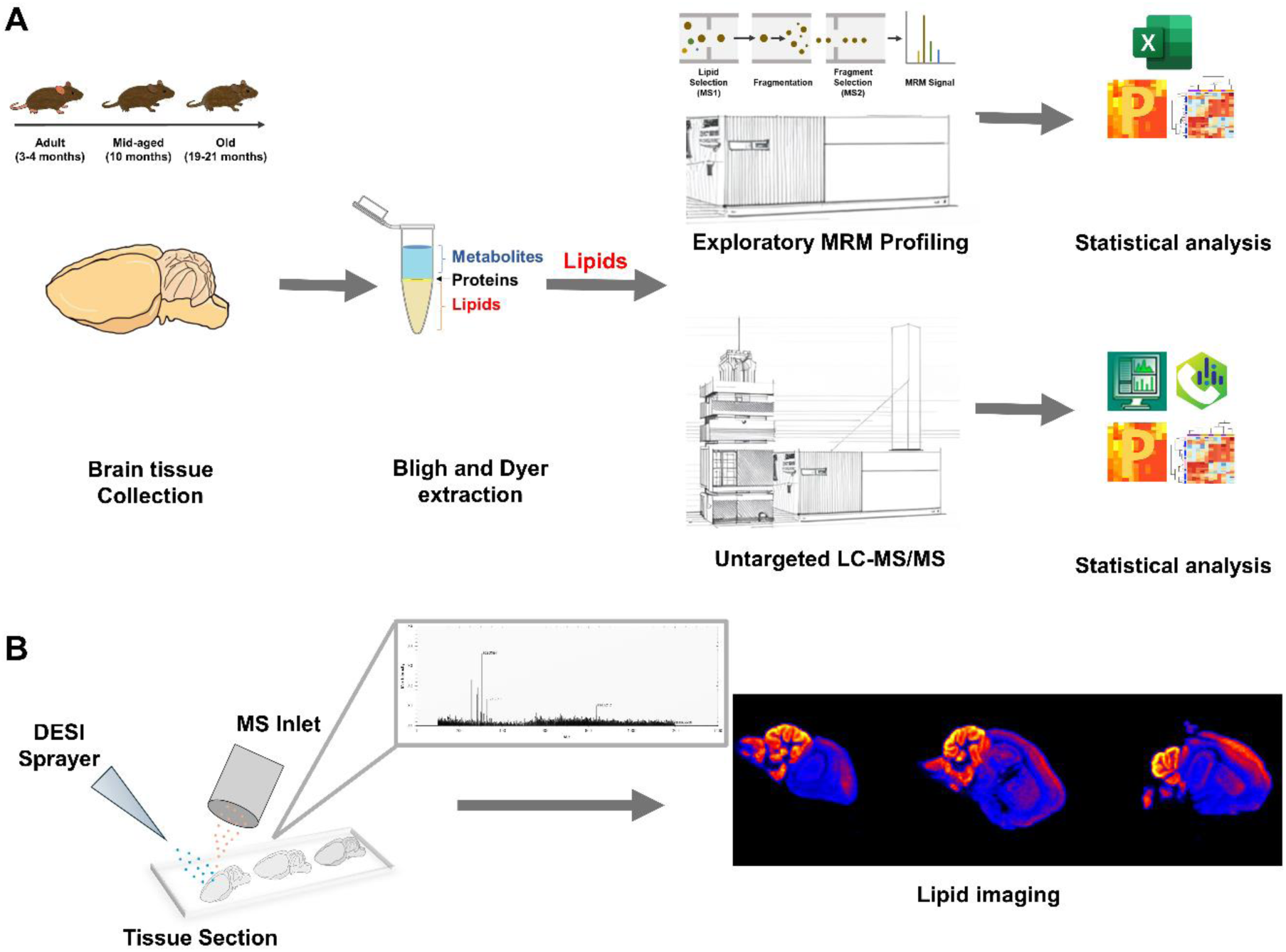
Experimental design. (A) Lipidomic analysis of mice’s brain samples. Whole brain tissues from three different age groups of mice were collected, homogenized, and lipids were extracted by the Bligh & Dyer method. These lipids were subjected to shotgun MRM profiling and untargeted LC-MS/MS profiling experiments. The shotgun MRM profiling data was analyzed using an internal script for data filtering. The LC-MS/MS data were qualitatively analyzed using Agilent MassHunter followed by data processing in MS-DIAL. Statistical analysis was performed using Perseus and OriginPro 2022 for both the MRM and LC-MS/MS data. (**B**) For DESI-MS imaging, cryomicrotome-sectioned brain tissues from the three age groups were subjected to ambient ionization using DESI spray, and data were collected at a resolution of 150 µm pixel resolution.

The 3246 MRM transition list was based on known precursor ion (Prec) and neutral loss (NL) scans that generate diagnostic fragments of classes or fatty acid diagnostic fragments [25, 31], which are specific to each lipid class of lipids (Figure 2A). Therefore, all lipid species are interrogated with MRM scans and the MRM scans were based on information from the Lipid Maps database combined with class- or structure-related product ions. The sample was continuously introduced into the ion source via flow injection using an autosampler, where lipids were ionized in positive or negative ion mode. PC, lysoPC, SM, PS, PE and lysoPE were analyzed as singly charged positive ions [M + H] +, and PG, PI, and PS were analyzed as singly charged ammonium adducts [M+NH4]+ [24].

**Figure 2:**
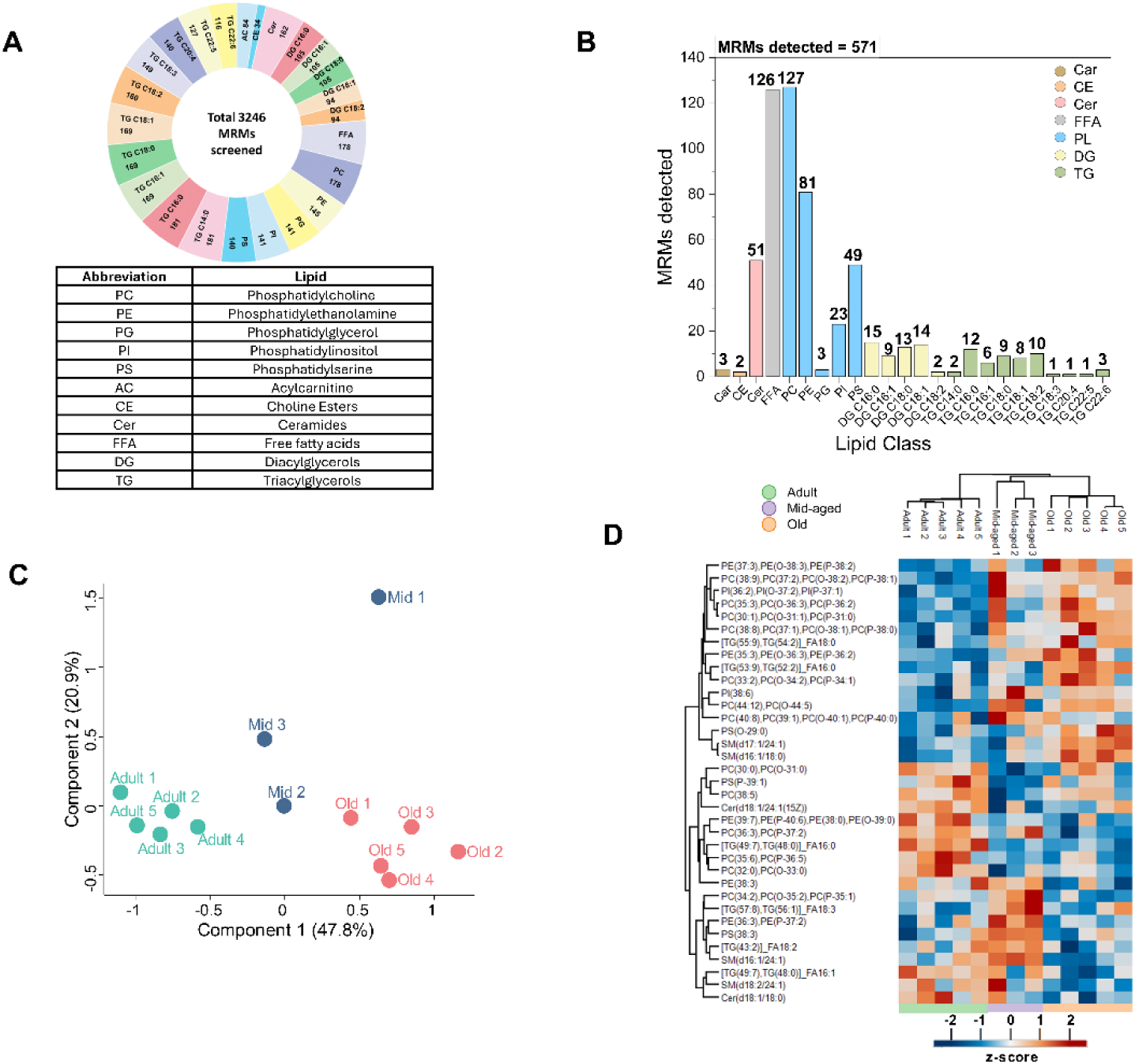
Results of MRM profiling. **(A)** For the MRM profiling, samples were screened for 3246 MRMs that span 11 lipid classes and the number of MRMs screened for each class is shown. **(B)** The number of MRMs detected in each lipid class. A total of 576 of the 3246 MRMs were successfully identified. **(C)** Principal component analysis (PCA) of samples from three different age groups of mice. **(D)** Hierarchical clustering and heat map of significantly changing MRMs. Cluster analysis was performed by Euclidean method for both rows and columns. A total of 35 MRMs were identified as significantly changing between the three age groups.

Each MRM used and the corresponding lipid attributed from the characteristic features of the MRM scans are presented in Supplementary Table S1. Data on MRM profile was collected for 13 brain samples representing five adult mice (3 to 4 months), three middle–aged mice (10 months), and five old mice (19–20 months). The intensity of each MRM was compared with a blank sample (injection solvent), and MRMs that showed ion intensities >30% of the ion intensity for the blank were retained for further analysis, resulting in 571 MRMs considered detected (Supplementary Table 1, Figure 2B) from a total of 3,246 MRMs that were screened. Of the 571 MRMs detected, PC was the most represented (22.05%), followed by FFA (21.8%), PE (14%), Ceramides (8.8), and PS (8.5%) (Figure 2B). As fatty acids could not be verified by further derivatization, they were not considered for biological interpretation from the MRM data, resulting in a total of 445 MRM that were considered for statistical analysis. The PCA plot (Figure 2C) clearly shows how replicates within the same age group cluster together and separate from other age groups, indicating similarity in lipid profiles within the same age group.

These 445 MRMs were subjected to multivariate analysis using ANOVA, resulting in the identification of 35 MRMs that were significantly different representing six lipid classes, including Cer, PC, PE, PI, PS and TG (Figure 2D, Supplementary Table 2). It is important to note that despite significant changes in these MRMs, their abundance distributions were variable. PE (0-38:3), PC(37:2), TG(54:2) and PI(38:6) showed a steady increase, while PC(38:5), PS(P-39:1),

Cer(d18:1/24:1) showed a gradual decrease with age (Figure 2D, Supplementary Table 2). TG(56:1), FA(18:3), PE(36:3), PS(38:3), PC(34:2) were most abundant in middle-aged mice compared to other age groups.

### Untargeted LC-MS/MS profiling of the aging mouse brain lipidome

In untargeted LC-MS/MS analysis (Figure 1), we identified 1,223 precursor ion (MS1) signals, including 587 ions detected in the positive ESI mode and 636 ions detected in the negative ESI mode, after filtering out the blanks (Supplementary Table 3, 4, 5, 6). Of these ions, 245 MS1 ions in positive mode and 154 MS1 ions in negative mode had MS / MS characteristics, from which 218 ions (150 in positive mode and 68 in negative mode) were matched to the MassBank of North America (MONA) reference database with their acquired MS/MS spectra (Figure 3A, 3B, Supplementary Figure 1A, 1B, Supplementary Table 3, 5). To expand the identification of lipids, we also searched LC-MS/MS data using National Institute of Standards and Technology (NIST) Mass Spectral Library Altas (Supplementary Table 4, 6). This search led to the identification of 106 and 93 reference-matched lipids in the positive and negative ion modes, respectively. By combining the two search results, we were able to identify a total of 153 lipids in positive ion modes and 109 lipids in negative ion modes that could be matched to reference databases, comprising ∼ 22% of the total acquired spectra excluding blanks (Figure 3A, 3B). These reference-matched lipid species were used for biological interpretation. Among the remaining spectra, most were either suggested (without MS/MS match) or unknown (Supplementary Figure 1A, B, Supplementary Tables 3, 4, 5, 6). It is important to note that more than 78% of the spectra were still not mapped to any reference database, highlighting the need to expand existing MS/MS reference library databases for comprehensive lipidome analysis.

**Figure 3:**
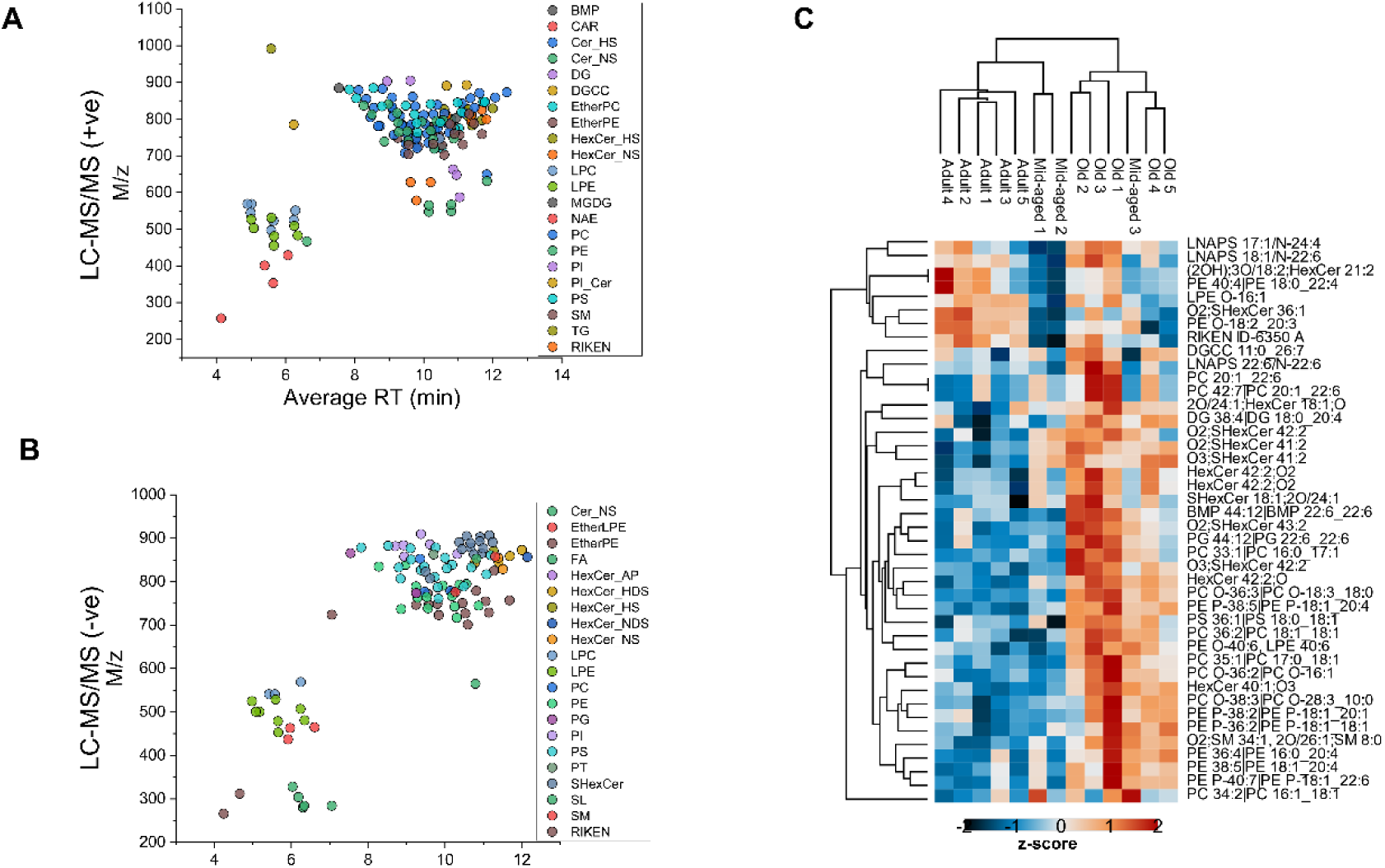
Results of untargeted lipidomics. **(A)** Scatterplot showing the 149 lipids matched to the MSP spectral database identified in the positive ion mode. **(B)** Scatterplot showing the 68 lipids matched to the MSP spectral database identified in negative-ion mode. Lipoproteins from different classes are color-coded. **(C)** Hierarchical clustering and heat map of significantly changing lipids identified from untargeted LC-MS/MS analysis. Among the significant lipids, a total of 40 lipids were reference matched from both positive and negative ion modes. Hierarchical clustering was performed using the Euclidean method for both rows and columns.

We performed statistical analysis on all 1,223 lipids identified in the MS1 dataset and identified 116 lipid characteristics that change in positive ion mode and 70 lipids that change in negative ion mode (Supplementary Tables 7, 8). A total of 40 significant lipids were reference matched (Supplementary Table 9) and most of these 40 significant lipids were more abundant in older mice than in adult or middle-aged mice. The PCA plot shows how the replicates of the different age groups are clustered separately, indicating differences in their lipid profiles (Supplementary Figure 1C, 1D). These significantly different lipids belong to classes that include phospholipids, ceramides, and diacylglycerols, suggesting that aging affects various classes of lipids and metabolic pathways.

Most of the lipids identified by untargeted analysis were also represented in the MRM profiling results, indicating a good overlap between the two methods, enhancing the reliability and robustness of the findings. Glycerophospholipids, PC and PE, which are the main components of cell membranes and control the membrane anchoring of proteins in brain cells [32, 33], were notably the most predominant lipid species identified in old mice (Figure 3C, Supplementary Table 9). PCs such as O-36:2, O-36:3 and O-38:3, as well as N-acyl-lysophosphatidylserines (LNAPS) such as LNAPS 17:1/N-24:4 and PS (36:1), were all higher in older mice compared to adult and middle-aged mice (Figure 3C, Supplementary Table 9). PS lipids with a high number of unsaturated fatty acids, such as DHA in PS (40:6), on the other hand, showed a trend of reduced levels in older mice; however, the data do not provide enough strong evidence to confirm that this reduction is not due to random variation.

To reveal any patterns or differences between the brains of adult and old-aged mice, we used volcano plots to represent the most highly changing lipids (Supplementary Figure 2A, 2B, 2C). The volcano plots for the changes in lipids between old and adult mice identified by the MRM analysis, untargeted LC-MS/MS (positive) and LC-MS/MS (negative) are shown in Supplementary Figures 2A, 2B, and 2C, respectively. It is evident from the distributions of these lipids in these graphs that the proportion of lipids that are up-regulated with age is higher than those that are down-regulated with age. From the MRM profiling data, phosphatidylserines such as PS (40:6), PS (42:1) and DG (42:6)_FA16:0 are the most highly downregulated with age in the brains of older mice, while SM (d16:1/18:0), PC (30:1), PC (35:3) and SM (d17:1/24:1) are the most highly upregulated lipids in the brains of old mice compared to the adult (Supplementary Figure 2A). Similarly, from the untargeted analyzes, SHexCer(36:1) and PE(O-38:5) are the most highly downregulated lipids in the brains of old mice, while PC(O-36:3), PC(O-38:3), DG(38:4), HexCer(40:1), SHexCer(41:2) and SHexCer(42:2) are the most highly upregulated lipids in the brains of old mice (Supplementary Figure 2B, 2C).

Ceramides and sphingolipids were identified in both MRM and non-targeted LC-MS/MS profiling, and their levels generally increased with age (Supplementary Tables 2, 9). Elevated levels of ceramides have previously been reported in AD and PD [34]. Cer(d18:1/18:0) and SM(d16:1/24:1) decreased in the brain with age (Supplementary Figure 2D, Supplementary Table 2). HexCers, the most abundant neutral glycosphingolipids in the myelin sheath, showed increased levels in the brains of old mice (Figure 2D, Supplementary Figure 2E). Furthermore, SM 34:1;O_2_, a sphingomyelin (ceramide phosphocholine) also increased with age.

Sulfatides or sulfoglycerophospholipids, which carry a sulfate ester group, are particularly concentrated in the substantia nigra region of the mouse brain [35]. Sulfatides with two double bonds, such as SHexCer 42:2;O_2_, SHexCer 41:2;O_2_, and SHexCer 43:2;O_2_, were more than three times higher in abundance in old mice compared to adult mice (Figure 3C, Supplementary Table 9). Specifically, sulfatides with only one double bond, such as SHexCer 36:1;O_2_, decreased in abundance with age (Figure 5B-i). On the contrary, SHexCer 42:2;O_3_, and HexCer 40:1;O_3_ were 5.3 and 3.6 times more abundant in the brains of old mice than in adult mice (Figure 5B-ii). These findings align with the sulfatide levels patterns observed in renal cell carcinoma, providing a link between altered sulfatide levels, aging, and cancer [36].

### DESI imaging analysis and comparison with MRM profiling and untargeted LC-MS/MS

DESI-MSI data were acquired in both negative and positive ion modes and the corresponding spectra were viewed using the MassLynx Spectra viewer. Given the functionally distinct regions of the brain, we performed segmentation analysis to understand how lipids are clustered in different areas of the brain. Using the uniform manifold approximation project (UMAP) classification for nonlinear dimensionality reduction for segmentation analysis, the generation of clusters from the UMAP space simplified data visualization by clustering the different lipid profiles in the UMAP space (Figure 4A). In the negative ion mode, six clusters were identified, clearly separating lipids in the cerebellum, cerebral cortex, pons and medulla, and other regions (Figure 4A). Cluster 1 (blue) and cluster 2 (orange) represent the sections of cerebellum and pons + medulla (undifferentiated), respectively. We can infer that their lipid profiles differed greatly from the rest of the brain, from the distances in the UMAP coordinate 1 (Figure 4A, Supplementary Figure 3A, 3B). Hierarchical clustering of 215 DESI-MSI features after removing background ions showed four major groups (Supplementary Figure 3C, Supplementary Table 11). Lipids in group 1 were high in the pons and medullary region and low in the cerebellar region, even though these structures are closely located in the brain (Figure 4A, Supplementary Figure 3). The lipids in group 2 were particularly high in the hypothalamus, basal ganglia, and midbrain. Sixty-six lipids, including [FA (20:4)-H]-, [PC (34:1)+Cl]-, [PS(40:6)-H]- and [PI(38:4)-H] were present in group 2. Group 3, consisting of [FA (18:1)-H]- and sulfatides like [SHexCer 36:1;O2-H]-, [SHexCer(42:2)O2-H]- and [SHexCer 42:2;O3-H]. Group 4 consisted of lipids like [FA (22:6)-H]-, [PE (40:6)-H]- and [PG (44:12)-H]-. Lipids in cluster 4 and 5 showed an increase with age, while cluster 1 showed decrease with age. Lipids in cluster 6 that occupied most of the regions of the brain showed a gradual shift in localization into the midbrain region with aging (Figure 4A, Supplementary Figure 3C, Supplementary Table 11).

**Figure 4:**
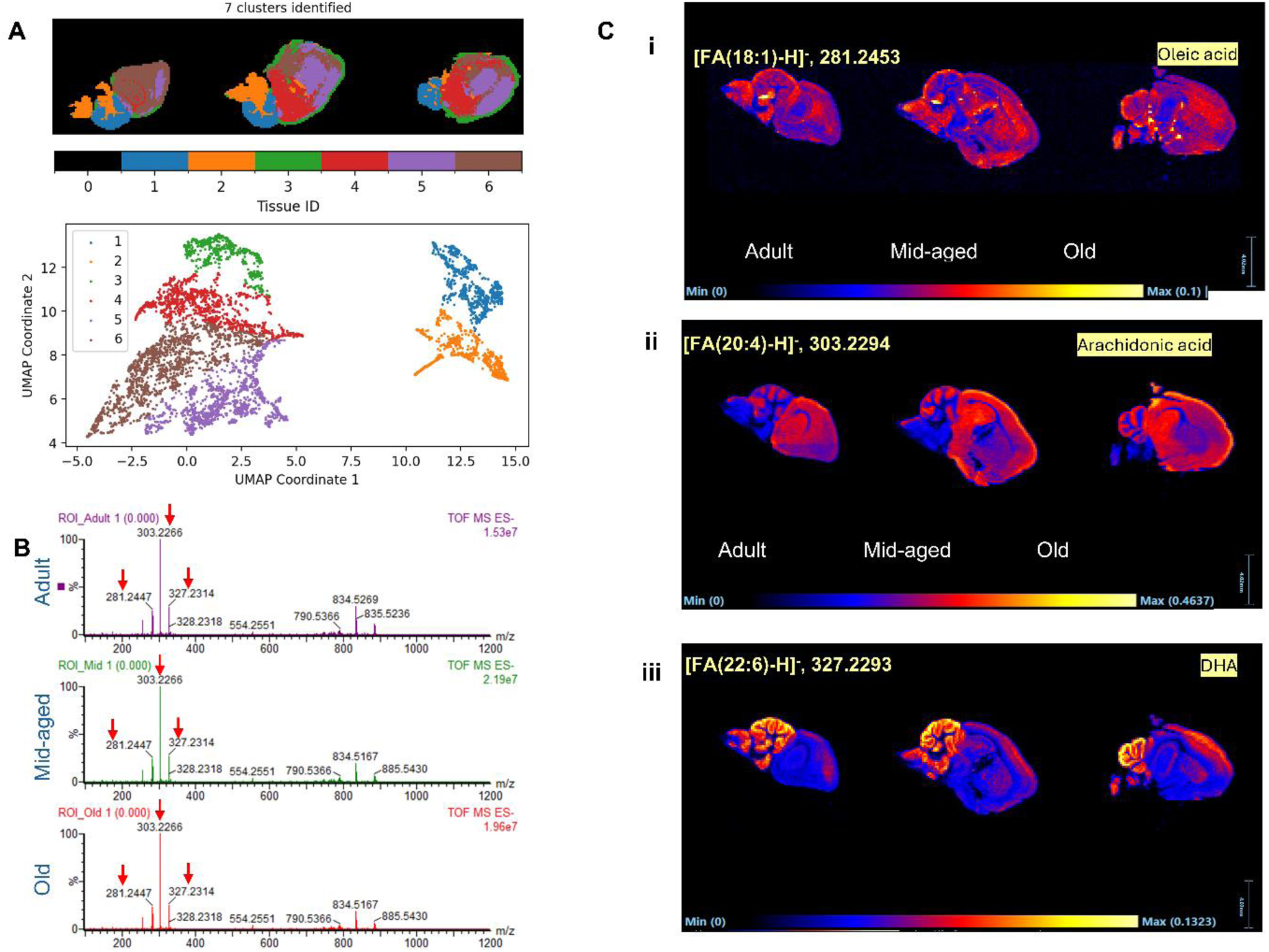
DESI-MSI of the aging mouse brain. **(A)** Segmentation and UMAP analysis of the DESI MSI (negative) data. Based on spectral characteristics, the regions in the brain could be divided into six groups (1-6) as shown. **(B)** DESI MSI (negative) spectra of the aging brain. The data was divided into three ROIs that refer to the three whole aging brains and the spectra of those were visualized. Fatty acids such as oleic acid (281.2453), arachidonic acid (303.2294), and DHA (327.2293) were the highest peaks that indicated their abundance in the brains. Apart from these, other abundant peaks identified in the negative ion mode DESI-MSI are m/z 834.5212 [PS (40:6)-H]-, m/z 790.5330 [PE (40:6)-H]-, and m/z 885.5419 (PI (38:4). **(C)** DESI-MSI of (i) oleic acid, (ii) arachidonic acid and (iii) DHA shows the spatial distributions of these lipids. Oleic acid is distributed throughout the brain, arachidonic acid is high in gray matter, and DHA is very high in the cerebellar regions and apparently spreads to the cortical regions with aging.

**Figure 5:**
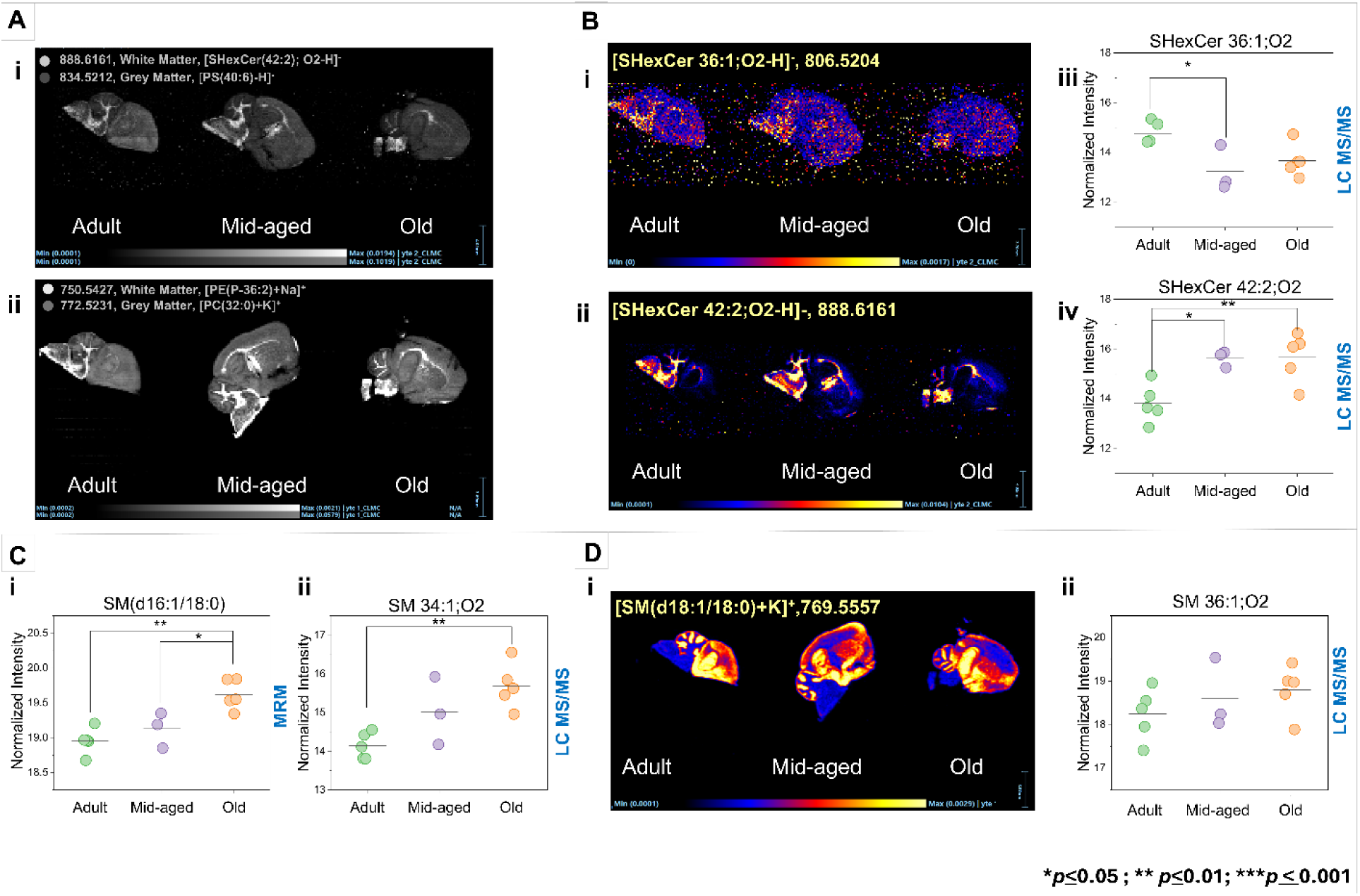
DESI-MSI and scatterplots of representative lipids across the brain. **(A)** Characteristic grey and white matter lipids identified in **(i)** DESI MSI (ve): m / z 888.6161 [SHexCer (42:2); O2-H]^-^ in white matter and *m/z* 834.5212 [PS (40:6)-H]^-^ in gray matter. **(ii)** DESI MSI (+ve): 750.5427 [PE (P-36:2)+Na]^+^ in white matter, *m/z* 772.5231 [PC (32) + K] + in grey matter. **(B)** DESI-MSI and corresponding LC-MS/MS data of sulfatides depicting changes in aging mouse brains. (i) *m/z* 806.5204 [SHexCer 36:1;O2-H]^-^ shows a decreasing pattern with age. **(ii)** 888.6161 [SHexCer 42:2;O2-H]^-^ increased with age. (iii) Scatter plot of LC-MS / MS SHexCer 36:1;O2 showing decease and (iv) Scatter plot of LC-MS/MS SHexCer 42:2;O2 showing increase with age. **(C).** SM(34:1);O2 identified in both MRM profiling and untargeted LC-MS/MS. SM(34:1);O2 was identified by both methods and showed an increasing pattern with age. **(D)** DESI MSI of [SM(d18:1/18:0)+K]^+^, an abundant sphingolipid showing increased abundances with age.

Fatty acids such as oleic acid (*m/z* 281.2453), AA (*m/z* 303.2294), and DHA (*m/z* 327.2293) were the highest peaks (Figures 4B, 4C). Oleic acid was distributed throughout the brain in all three ages, while AA is high in gray matter, and DHA was very high in the cerebellar regions and apparently spread to the cortical regions with age. AA [FA (20:4) - H] and DHA [FA (22:6) - H] are the two most abundant polyunsaturated fatty acids in the brain that comprise 25% of brain fatty acids (Figure 4B, 4C) [37]. These were high in the cerebellum and the cerebral cortex, as previously reported. DHA, an omega-3 fatty acid, is an important component of the neural membrane and has been well studied over the years for its contribution to cognitive health during aging in the human brain [38–40]. Both lipids were also detected in our shotgun MRM-MS and untargeted LC-MS/MS data as well (Supplementary Table 1,5). The characteristic lipid ions in the positive and negative ion mode for gray and white matter (Figure 5A, Supplementary Figure 4A, 4B, Supplementary Table 12) were used to design the ROIs for these regions [28].

Figure 5B shows the sulfatides that showed interesting patterns and gave us similar observations from the untargeted LC-MS/MS and DESI-MSI. Sulfatides such as SHexCer (42:2);O2 (Figure 5A-i), that are characteristic lipids of the white matter were visualized with DESI-MSI and increased with age. We verify several lipids that change in the brains of aging mice and visualize their spatial distributions using DESI-MSI (Figure 6A). For example, [PC (34:2)^+^H]^+^, *m/z* 758.5546 was high in middle-aged mice in both MRM profiling (Figure 6A-i) and untargeted LC-MS/MS analyses (Figure 6A-ii). From the sagittal section, it was evident that it is distributed throughout the brain, especially in the cortical regions, except for the pons and medulla region (Figure 6A-i). It was interesting to note the increasing pattern of this lipid in the cerebellum with age, which can be confirmed by future studies with statistics from these imaging data.

**Figure 6:**
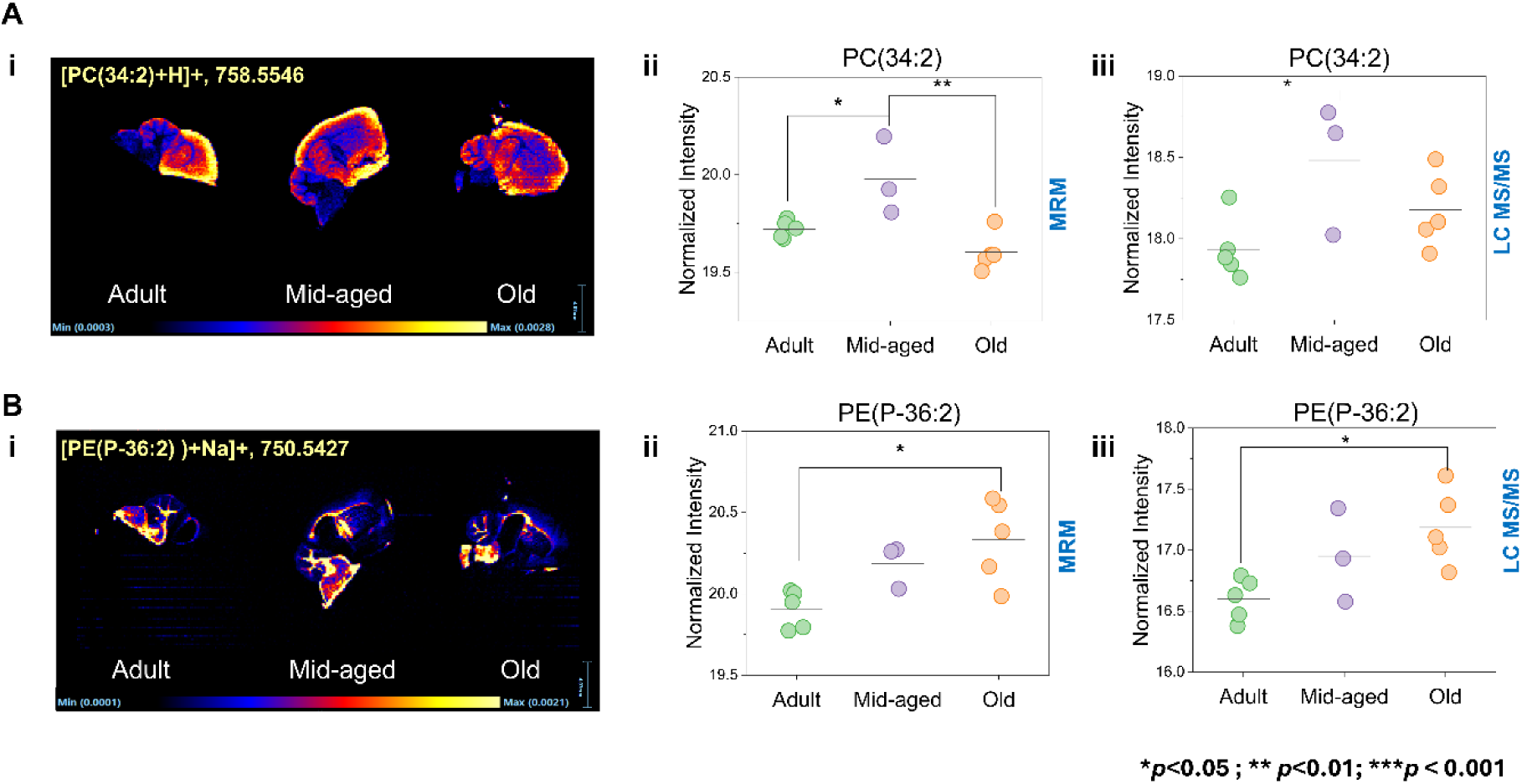
Examples of lipids identified in all three methods show an increase in abundance with aging. **(A)** PC (34:2), phosphatidylcholine was identified in all three analyzes and its spatial distribution is seen in the cortical regions, as well as in the midbrain and basal ganglia. **(B)** PE(P-36:2), a plasmalogen was identified in the three analyzes and showed an increase in abundance with age.

Plasmalogen, [PE(P-36: 2) + Na] + or [PE(O-36: 3], m / z 750.5427 increased with age in the brain from DESI MSI (Figure 6B-i), MRM profiling (Figure 6B ii) and untargeted LC-MS/MS data (Figure 6B-iii). It is in the white matter of the brain, as clearly indicated in the DESI MSI images (Figure 5A-ii; Figure 6B-ii). PC(O-36:3), another plasmalogen glycerophosphocholine was also detected in all three analyses and increasing in the aging mice brain (Supplementary Figure 5A). Phosphatidylglycerol PG(44:12), that was high in the cerebellar region and has also been reported earlier as a signature lipid of the mouse hippocampal region [41], was detected in LC-MS/MS analysis also and increasing in the brain with age (Supplementary Figure 5B). Phosphatidylinositol PI(38:4; *m/z* 885.4519) decreased in the brain with aging (Figure 7A), and although not statistically significant, they clearly decreased in the brain of aging mice, in MRM profiling and untargeted LC-MS/MS data. Previously, glycerophosphatidylinositols such as PI (38:4) have been reported to decrease in AD mouse models and may be related to impairments in memory and cognition [42]. On the other hand, PIs like PI (38:6) showed an increase with age in both DESI and MRM data (Supplementary Figure 5C). PS (40:6) is high in brain gray matter (Figure 5A-i, Supplementary Figure 4A), which is rich in glia and mainly unmyelinated neurons, and has a high content of DHA and stearic acid. It is a characteristic lipid in the gray matter of the brain [28] identified as a negative ion [PS(40:6)-H] in brain DESI-MSI (Figure 7A-i). The T tests between the old and adult age groups revealed that this lipid is significantly different (p-value=0.015), specifically it was shown to decrease 0.8 times in the old brain compared to the adult brain. Although not statistically significant, the decreasing pattern was clear in the MRM profiling (Figure 7B-ii) and untargeted data (Figure 7B1-iii). DESI-MSI also revealed a decreasing distribution of this lipid with age.

**Figure 7:**
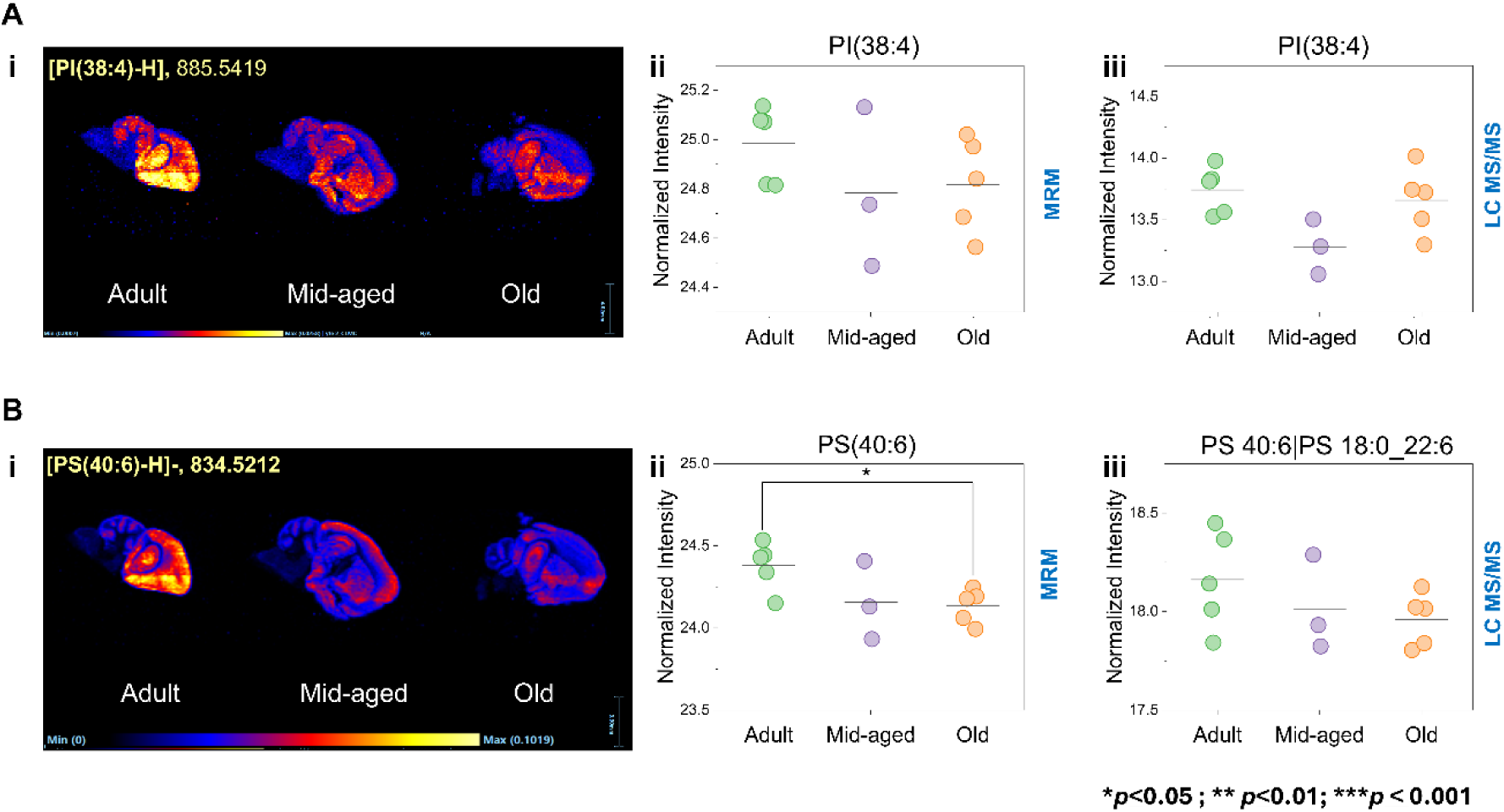
Examples of lipids identified in all three analyzes showing decreasing abundances with age. **(A)** PI (38:4), a phosphatidylinositol was identified in the three analyzes and showed decreasing abundances with age in both MRM profiling (ii) and untargeted LC-MS/MS data(iii). **(B)** PS (40:6), phosphatidylserine was decreasing in the brain with age, as seen in DESI MSI(i), MRM profiling(ii) and untargeted LC-MS/MS (iii) data.

### Interrogation with the proteomics data

Our analysis showed a clear decrease in DHA rich lipids, such as PS (40:6), in the brains of aging mice (Figure 7B). Polyunsaturated fatty acids (PUFA) such as DHA also declined in the orbitofrontal cortex, reducing gray matter and causing neuronal shrinkage. This decline is accompanied by increased levels of enzymes such as desaturases and elongases [43]. Although we did not identify these enzymes, our proteomic analysis showed that enzymes involved in fatty acid elongation, such as very long chain (3R) -3-hydroxyacyl-CoA dehydratase 3 (Hacd3), were five times higher in the brains of older mice compared to adults (Supplementary Table 10, Supplementary Figure 6A), suggesting a compensatory increase in these enzymes as specific lipids decrease with age. On the contrary, enzymes crucial for mitochondrial fatty acid beta oxidation, such as medium and very long chain specific mitochondrial acyl-CoA dehydrogenase (Acadm, Acadvl), decreased significantly with age (Supplementary Table 10, Supplementary Figure 6A) corroborating previous results [44]. Enzymes such as medium-chain and very long-chain specific mitochondrial acyl-CoA dehydrogenase decreased significantly with age, as indicated by our previously published proteomics data. These are important enzymes that catalyze mitochondrial fatty acid beta-oxidation. Dysregulation of their pathways has been reported in neurodegenerative diseases and aging [44].

By mapping the lipids identified in this study (Supplementary Figure 7) and correlating them with proteins identified from our recent proteomic work using the same tissues (https://doi.org/10.1016/j.mcpro.2024.100819), we identified proteins such as alpha-1,3/1,6-mannosyltransferase (Alg2), acid ceramidase (Asah1), arylsulfatases (ArsA, ArsB) and phosphatidylinositol 5-phosphate 4-kinases (Pip4k2a) (Supplementary Figure 6, Supplementary Table 10). While Alg2, Asah1, and Pip4k2a showed changes in total protein levels (Supplementary Figure 6A), sphingomyelin phosphodiesterase 3 (Smpd3) and phosphatidylinositol 4-kinase (Pi4k) only changed at the phosphoprotein level (Supplementary Figure 6B, with the alpha (Pi4ka) subunit increasing with age and the beta subunit (Pi4kb) decreasing with age (Supplementary Table 10). This highlights the complex interplay between lipids and proteins and protein regulation by modifications such as phosphorylation.

## DISCUSSION

Brain functions can vary greatly between different regions of the brain. Lipids are a crucial part of the complex information processing system of the brain. The study of these lipids using mass spectrometry (MS) is essential to understand how our biology changes as we age. Lipids play a critical role in cell structures, energy storage, and cell signaling. MS has emerged as a powerful tool that helps scientists to accurately identify, measure, and map the spatial distribution of many different lipid types in cells or tissues. This gives us detailed information on how lipids are organized and altered as we age. Changes in the composition and function of lipids are associated with many age-related conditions, such as heart disease, brain disorders such as AD or PD, and metabolic syndromes such as diabetes. By using three different MS-based analytical methods, we aim to understand the complex patterns of lipids associated with aging. We also sought to find specific lipids that could act as biomarkers for age-related diseases. This knowledge could ultimately lead to the development of treatment strategies to adjust lipid pathways, promoting healthier aging and improved quality of life throughout aging.

Aging increases oxidative stress and inflammation in the brain, impacting the regulation and function of lipids and other biomolecules. Understanding how the composition, structure, function, and spatial organization of lipids change with age in the brain is crucial to revealing the mechanisms underlying aging and age-related brain function. Unfortunately, this has been challenging because of the limitations of analytical methods. Recent advances in MS-based techniques now allow a comprehensive and precise analysis for the detection of lipids in diverse samples. Typical MS-based methods include MRM-profiling by direct infusion, global and targeted analysis based on LC-MS/MS, and MS imaging. These powerful techniques, coupled with dedicated extraction protocols, have demonstrated impressive analytical performance in terms of sensitivity, reproducibility, linearity, and resolution of isomeric lipid species from limited sample amounts. In this study, we successfully combined these three analytical methods to reveal changes in lipid composition and distribution in the brains of aging mice.

Aging is the hallmark of neurological disorders [45, 46] Therefore, identifying lipids as molecular signatures of aging may provide clues to the onset and progression of AD and PD [47]. Our integrated approach enabled the analysis of diverse lipids at the same time, including apolar lipids such as triglycerides and cholesterol esters together with polar lipids such as glycerophospholipids and sphingolipids such as sphingomyelin and ceramide. The compatibility of the methods enabled accurate identification and quantification of lipids, potentially enhancing our insights into the functions of various lipid classes and individual lipid species. Our results suggest an increasing susceptibility of the aging brain to neurodegenerative diseases. Triglycerides, TG 52: 2 increased and TG 48: 0 decreased with age, suggesting a protective role against cognitive decline. fatty acids (FA 24:5) and sphingomyelins (SM 22: 0 and SM 24:1) were among the most abundant lipids in the brain of old mice. The observed decrease in DHA-rich lipids, such as PS (40:6), aligns with previous findings linking the decline of polyunsaturated fatty acids (PUFAs) such as DHA to reductions in gray matter and neuronal shrinkage in the aging brain [38].

Neurodegeneration is an age-related pathological condition in which the nervous system and/or individual neurons lose structure, function, or both leading to progressive neural degeneration and cognitive and motor decline. Ceramides have been shown to progressively increase with aging in both cultured cells and whole brain samples [48]. C6-ceramide, a cell-permeable analog of natural ceramides, has been shown to increase A production of Aβ in Chinese hamster ovary cells by promoting beta-cleavage of APP, a process that leads to the formation of Aβ [49]. Our data also emphasizes the important role of ceramides in aging. However, enzymes that are directly involved in the processing of APP into Aβ and the mechanisms underlying this process are not fully understood, warranting further investigation. Sphingolipids such as sphingomyelin (SM), gangliosides, and cerebrosides are all derived from ceramides (N-acetylsphingosine) [50] and a connection between these ceramide derivatives and AD has been demonstrated [51]. Our results also corroborate these previous studies. Ceramides are the hydrophobic moiety of sphingolipids with long-chain fatty acylated bases that differ in their degree of saturation [52]. Sphingolipids are associated with maintaining brain functions via neurogenesis, and synaptogenesis [53]. Elevated levels of sphingomyelins, such as SM (d16:1/24:1) with age (Figure 5 & Supplementary Figure 2) suggest a compensatory role of these specific SM in maintaining brain function and the potential to contribute to age-related changes in brain function. Integrating lipidomic results with our recent proteomic results using the same tissues (DOI: http://doi.org/10.1016/j.mcpro.2024.100819), we found that acid ceramidase (Asah1), an enzyme responsible for the conversion of ceramides (N-Acyl-sphingosine) to sphingosine (KEGG reaction ID: R01494), increased in older mice (Supplementary Figure 2D). This is in agreement with the decrease in Cer (d18:1/18:0) in the brains of older mice determined by MRM profiling and the increase in sphingomyelins such as SM (34:1;O_2_) in both MRM profiling and untargeted LC-MS/MS. Furthermore, elevated levels of glycosphingolipid sulfates in older brains highlight their importance in signaling and brain function. Sulfatides show varied expression patterns with age, indicating their complex roles in aging and disease [54]. Sphingolipids of synaptic membranes interact with neurotransmitter receptors and regulate their activity [55]. A high level of SM occurs in the white matter of the brain as the primary component of the myelin membrane [56]. Gangliosides are abundant in the CNS, including the brain, and are associated with cell signaling and neuroprotection [57]. Although there have been many reports of the involvement of SMs in the brain, their identity remains elusive. Our study fills some gaps in our knowledge about the identification of specific sphingolipids and links them to aging. Although the involvement of ceramides and their derivatives in aging is clear [58], the precise mechanisms and biological significance of ceramide diversity in aging and neurodegenerative diseases require further investigation.

PC was identified as the most abundant phospholipid, followed by PEs, PSs, and PIs (Figure 2B). These phospholipids were detected in both positive and negative ion modes, while PGs were identified only in the negative ion mode. Plasmalogen (P-36:2), a predominant ether-phosphatidylethanolamine lipid, was consistently detected in all three methods and demonstrated an increase in abundance with age (Figure 6B). Approximately 20% of the phospholipids in human tissue are plasmalogens, which are particularly enriched in the brain, heart, and immune cells [59]. Studies have indicated that plasmalogen accumulation is associated with brain development and maturation [60]. In murine models, plasmalogens have been shown to mitigate age-related synaptic defects and microglia-mediated inflammation [59]. Plasmalogen distribution is notably high in the caudal structures of the brain (Figure 6B). Furthermore, an increase in plasmalogen levels has been observed in extracellular vesicles derived from brain tissue in patients with Alzheimer’s disease (AD) patients [61]. In addition to the established role of plasmalogens in brain function and cognition, a decrease in PS (40:6) has also been documented in extracellular vesicles in AD models [61]. These findings underscore the importance of plasmalogens and other phospholipids in both normal brain physiology and the pathophysiology of neurodegenerative diseases such as AD.

In general, current results emphasize the importance of lipid regulation in the aging brain and its link to neurodegenerative diseases. As we age, the balance and composition of lipids in the brain can change significantly, which can contribute to the development and progression of conditions such as AD, PD, and other cognitive declines, as already suggested by other research [62–65]. These results highlight the need for further research into the specific ways in which lipid alterations affect brain function. Future research should focus on the functional implications of these lipid alterations, examining how they influence cellular processes, neuronal communication, and overall brain health. By exploring these mechanisms in greater detail, we can identify the critical points of intervention that can help maintain brain health as we age.

Furthermore, understanding the molecular mechanisms of lipid dysregulation due to aging and its impact on brain function could pave the way for novel therapeutic strategies. Such strategies might include the development of drugs that target specific lipids or their pathways, or lifestyle changes that support healthy lipid metabolism and could potentially slow or even prevent the onset of age-related neurological disorders. By focusing on qualitative, quantitative, and spatial differences in brain lipids, this study provides a better understanding of the critical role that lipids play in the aging process.

### Limitations of this study

This study contributes to our understanding of age-dependent changes in the brain lipidome. However, it is important to consider the specific limitations of the study for a complete understanding of the experimental context and interpretation of the results. First, shotgun MRM profiling and LC-MS / MS analyzes were performed using homogenized tissue samples. The brain is a complex organ, not only for its lipid composition but also for the spatial distribution of lipids to different brain regions that perform spatially resolved functions. Regions such as the cortex, hippocampus, and cerebellum differ markedly in their cellular architecture and function, and lipids are vital in these regions, which influence membrane fluidity, signaling pathways, and energy storage. The lack of spatially resolved and cell-type-specific lipidomic means that potentially crucial variations in lipid composition, abundances and metabolism that could reveal spatially resolved and cell-type-specific brain functions or pathologies remain unexplored.

Although we performed DESI-MS imaging, which allows us to determine spatial distribution of lipids in different brain regions, these data need to be validated using spatially resolved and cell-type-specific LC-MS/MS analysis. Second, this study only employed male mice. Many age-dependent changes in lipids are sex dependent [66]. Validating current results using female mice and investigating sex-dependent and independent variation of the brain lipidome during the aging process is necessary to generalize current results. Third, increasing the sample size in future studies could improve the robustness and precision of the data, allowing more robust conclusions to be drawn from such studies.

## Abbreviations

FA: fatty acids
BMP: bismonoacylglycerophosphate
Cer-HS: Ceramide hydroxyfatty acid sphingosine
Cer-NS: Ceramide nonhydroxyfatty acid sphingosine
LPC: lysophophatidylcholine
LPE: lysophosphatidylethanolamine
DG: diacylglycerol
ShexCer: sulfated hexosylceramides
HexCer: hexosylceramide
HexCer-AP: hexosylceramide alpha-hydroxy fatty acid-phytospingosine
HexCer-HS: hexosylceramide hydroxyfatty acid sphingosine
HexCer-NS: hexosylceramide nonhydroxyfatty acid sphingosine
HexCer-NDS: Hexosylceramide nonhydroxyfatty acid-dihydrosphingosine
HexCer-HDS: hexosylceramide hydroxyfatty acid-dihydrosphingosine
MGDG: monogalactosyldiacylglycerol
PC: Phosphatidylcholines
PS: phosphotidylserines
PI: Phosphatidylinositols
SM: Sphingomyelins
SHexCer: Sulfohexosylceramides
LNAPS: *N*-acyl-lysophosphatidylserine
Mid-aged: Middle-aged mice were referred to as “mid-aged” in the figures

## Supporting information

### Supplementary Figures

**Supplementary Figure 1.** Lipids detected in untargeted LC-MS/MS analysis.

**(A)** Scatterplot of 587 LC-MS/MS (+ve) identified lipids. This scatterplot represents the lipids identified in the aging mouse brain by using LC-MS/MS in positive ion mode, annotated by the MSP spectral database. Of the total, 214 lipids were unknown, 224 lipids either lacked MS/MS data or their MS/MS spectra could not be matched to the MSP spectral database, and 149 reference-matched lipids were identified with high confidence and considered for further biological interpretation.

**(B)** Scatterplot of 636 identified lipids identified by LC-MS / MS (-ve). This scatterplot displays the lipids identified in the aging mouse brain using LC-MS/MS in negative-ion mode. The data set comprised 134 unknown lipids, 434 lipids without MS/MS matches, and 70 lipids that could be matched with the MSP spectral database.

**(C)** PCA scores plot of LC-MS/MS (+ve) lipids. The PCA plots show a distinct clustering of adult, middle-aged and old mice, indicating differences in their lipid profiles identified by untargeted analysis.

**(D)** PCA scores plot of LC-MS/MS (-ve) lipids reveals separate clustering of adult, middle and old mice, indicating age-related variations in lipid profiles identified by untargeted analysis.

**Supplementary Figure 2:** Volcano plots of changes in lipids in the aged mouse brain compared to adults.

**(A)** MRM profiling. Volcano graph showing lipids with significant changes in the aged mouse brain compared to adults.

**(B)** Untargeted LC-MS/MS (+ve) analysis. Volcano plot depicting lipids with significant changes in the aged mouse brain compared to adults, identified using LC-MS/MS in positive ion mode.

**(C)** LC-MS/MS (-ve) analysis. Volcano plot illustrating lipids with significant changes in the aged mouse brain compared to adults, identified using LC-MS/MS in negative ion mode.

**(D)** Ceramides in MRM profiling. Ceramides identified by MRM profiling show a decrease in the aged mouse brain.

**(E)** Sphingomyelins and hexosylceramides. Examples of sphingomyelins and hexosylceramides identified in both MRM profiling and untargeted LC-MS/MS data that exhibit an increase in the aged mouse brain.

**Supplementary Figure 3:** Representative DESI-MSI spectra for tissue clusters 1, 2 obtained in DESI negative ion mode (A, B). The most abundant and notable peaks are highlighted.

(A) Heatmap of lipids corresponding to different regions of the brain identified by MSI segmentation and hierarchical clustering.

**Supplementary Figure 4: (A)** Notable lipids of grey and white matter identified in the DESI MSI analysis in positive ion mode. **(B)**. Notable lipids of grey and white matter identified in DESI MSI analysis in negative-ion mode. The peaks for these lipids were extracted and are indicated in **(C)** and **(D)**, respectively.

**Supplementary Figure 5:** Examples of phospholipids identified in more than one analysis. **(A)** PC(O-36:3), **(B)** PG(44:12), and **(C)** PI(38:6).

**Supplementary Figure 6:** Heat map of proteins significantly changing from proteomics data mapped from the lipidomics data in the brain of aging mice.

(A) Total proteins changing with the associated lipids such as ceramides, PIs, PEs.

(B) Phosphosites significantly changing with the lipids such as PI and Sphingomyelin changing in the aging brain.

**Supplementary Figure 7.** Pathway mapping of lipids and integration with previously published proteomics data in aging mice. Genes associated with identified lipids, either directly or indirectly, are represented as circles and color coded according to their significance in previously published proteomics data from the same set of mice. KEGG IDs for the significantly changing lipids were retrieved and analyzed using Metascape. The gene IDs were then matched to the proteomics data to elucidate the interconnected changes in lipids and proteins in the aging brain.

**(A)** Pathway mapping of ceramides, sphingomyelins, hexosylceramides, and sulfatides. This panel shows the pathway mapping for these specific lipid categories, highlighting their roles and interactions.

**(B)** Pathway mapping of phosphatidylinositols. This panel focuses on the mapping of the phosphatidylinositol pathway, detailing their interactions and significance.

## Supplementary data set

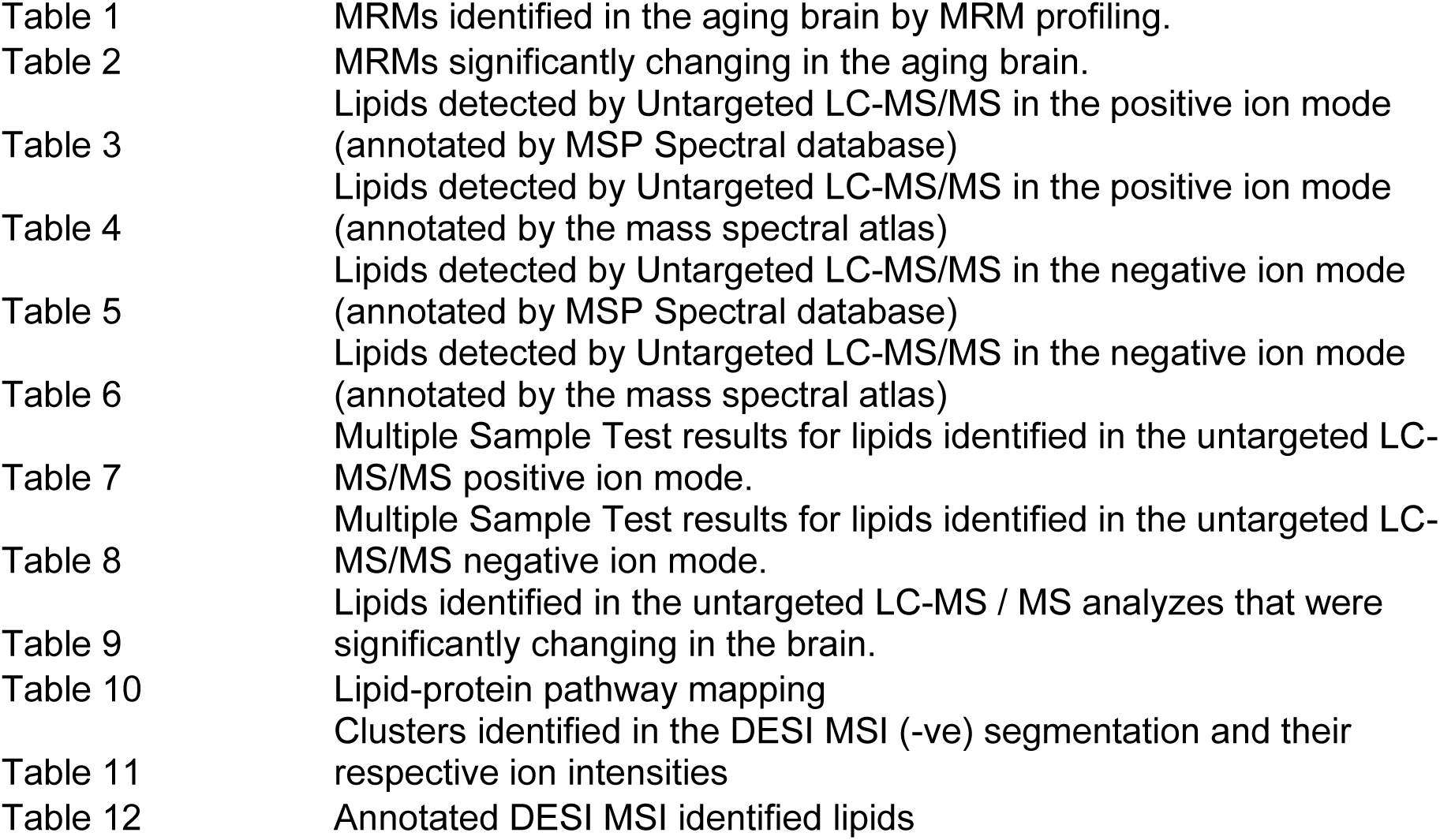

**Supplementary data set:**
|  |  |
| --- | --- |
| Table 1 | MRMs identified in the aging brain by MRM profiling. |
| Table 2 | MRMs significantly changing in the aging brain. |
| Table 3 | Lipids detected by Untargeted LC-MS/MS in the positive ion mode (annotated by MSP Spectral database) |
| Table 4 | Lipids detected by Untargeted LC-MS/MS in the positive ion mode (annotated by the mass spectral atlas) |
| Table 5 | Lipids detected by Untargeted LC-MS/MS in the negative ion mode (annotated by MSP Spectral database) |
| Table 6 | Lipids detected by Untargeted LC-MS/MS in the negative ion mode (annotated by the mass spectral atlas) |
| Table 7 | Multiple Sample Test results for lipids identified in the untargeted LC-MS/MS positive ion mode. |
| Table 8 | Multiple Sample Test results for lipids identified in the untargeted LC-MS/MS negative ion mode. |
| Table 9 | Lipids identified in the untargeted LC-MS / MS analyzes that were significantly changing in the brain. |
| Table 10 | Lipid-protein pathway mapping |
| Table 11 | Clusters identified in the DESI MSI (-ve) segmentation and their respective ion intensities |
| Table 12 | Annotated DESI MSI identified lipids |

## Acknowledgements

MS analysis was performed at the Bindley Bioscience Center, Proteomics, and Metabolite Profiling Facilities. We also thank the Histology Research Laboratory at Purdue University College of Veterinary Medicine for generous help with frozen tissue sections in a Leica CM1860 cryostat tissue sectioning for DESI-MSI analysis. This study was supported by funding from the Bindley Fellow Program (UKA), Indiana Clinical and Translational Science Institute (UKA), Purdue COVID disruption fund (UKA), Showalter Trust Grant (UKA. & AJS) and Purdue start-up fund (AJS). The Waters SYNAPT XS ion mobility time-of-flight MS used for DESI MSI experiments was supported by an NIH shared instrumentation S10 grant 1S10-OD026954-01, located at the Purdue Metabolites Profiling Facility, Bindley Bioscience Center.

## Contributions

PP: Conceptualization and experimental design, MS data collection, analysis and interpretation, and manuscript preparation. CRF: Experimental design, MS data collection, analysis and interpretation, and manuscript review. BRC: LC-MS/MS data acquisition, data analysis and interpretation, and revision of the draft manuscript. AJS: Housing and raising of mice, tissue collection, experimental design, data interpretation, manuscript review, and funding acquisition. UKA: Conceptualization and experimental design, project supervision, collection, analysis and interpretation of MS data, preparation of manuscripts, and acquisition of funds.

