## Supplementary Figures for "Multiplatform lipid analysis of the brain of aging mice by mass spectrometry"

### Slide 1
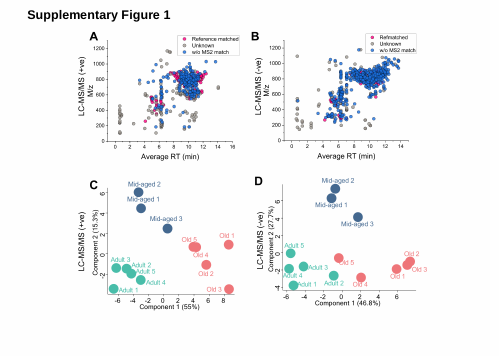

Supplementary Figure 1

### Slide 2
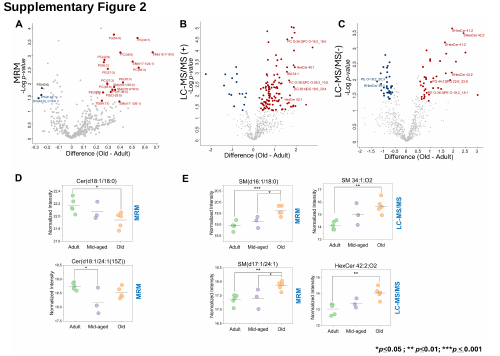

Supplementary Figure 2

### Slide 3
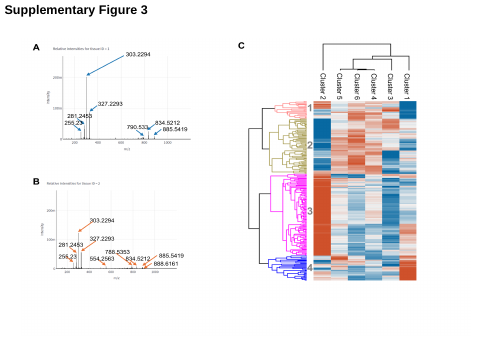

Supplementary Figure 3

### Slide 4
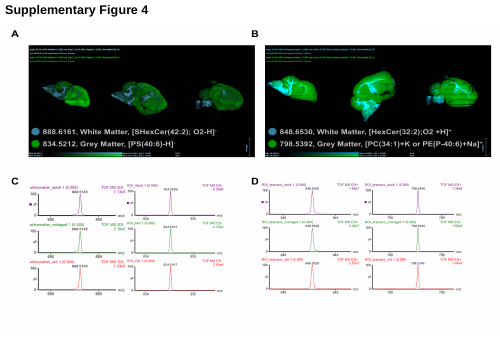

Supplementary Figure 4

### Slide 5
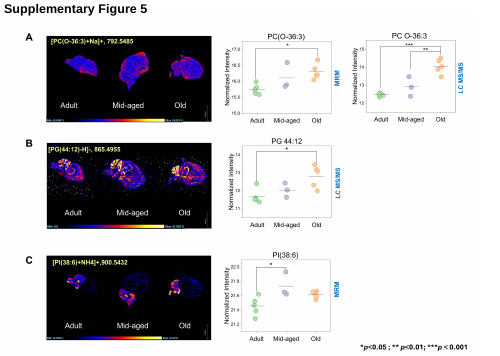

Supplementary Figure 5

### Slide 6
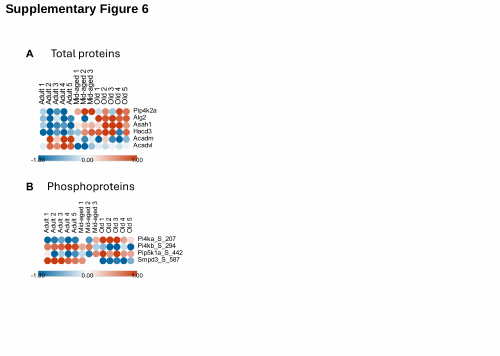

Supplementary Figure 6

### Slide 7
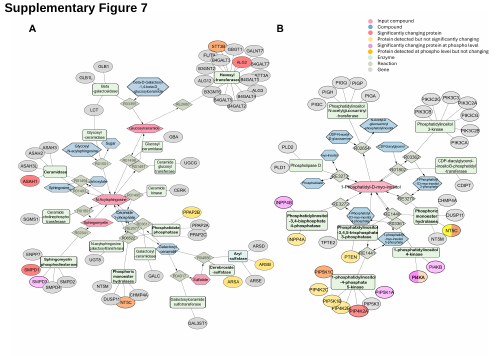

Supplementary Figure 7
